# Enhanced immunogenicity of a synthetic DNA vaccine expressing consensus SARS-CoV-2 Spike protein using needle-free immunization

**DOI:** 10.1101/2021.02.01.429219

**Authors:** Sawsan S Alamri, Khalid A Alluhaybi, Rowa Y Alhabbab, Abdullah Algaissi, Sarah Almahboub, Mohamed A Alfaleh, Turki S Abujamel, Wesam Abdulaal, M-Zaki ElAssouli, Rahaf Alharbi, Mazen Hassanain, Anwar M Hashem

## Abstract

The ongoing global pandemic of Coronavirus Disease 2019 (COVID-19) calls for an urgent development of effective and safe prophylactic and therapeutic measures. Severe acute respiratory syndrome coronavirus 2 (SARS-CoV-2) spike (S) glycoprotein is a major immunogenic and protective protein, and plays a crucial role in viral pathogenesis. In this study, we successfully constructed a synthetic codon-optimized DNA-based vaccine as a countermeasure against SARS-CoV-2; denoted as VIU-1005. The design was based on the synthesis of codon-optimized coding sequence for optimal mammalian expression of a consensus full-length S glycoprotein. The successful construction of the vaccine was confirmed by restriction digestion and sequencing, and the protein expression of the S protein was confirmed by western blot and immunofluorescence staining in mammalian cells. The immunogenicity of the vaccine was tested in two mouse models (BALB/c and C57BL/6J). Th1-skewed systemic S-specific IgG antibodies and neutralizing antibodies (nAbs) were significantly induced in both models four weeks post three injections with 100 μg of the VIU-1005 vaccine via intramuscular needle injection but not intradermal or subcutaneous routes. Importantly, such immunization induced long-lasting IgG response in mice that lasted for at least 6 months. Interestingly, using a needle-free system, we showed an enhanced immunogenicity of VIU-1005 in which lower doses such as 25-50 μg or less number of doses were able to elicit significantly high levels of Th1-biased systemic S-specific IgG antibodies and nAbs via intramuscular immunization compared to needle immunization. Compared to the intradermal needle injection which failed to induce any significant immune response, intradermal needle-free immunization elicited robust Th1-biased humoral response similar to that observed with intramuscular immunization. Furthermore, immunization with VIU-1005 induced potent S-specific cellular response as demonstrated by the significantly high levels of IFN-γ, TNF and IL-2 cytokines production in memory CD8^+^ and CD4^+^ T cells in BALB/c mice. Together, our results demonstrate that the synthetic VIU-1005 candidate DNA vaccine is highly immunogenic and capable of inducing long-lasting and Th1-skewed immune response in mice. Furthermore, we show that the use of needle-free system could enhance the immunogenicity and minimize doses needed to induce protective immunity in mice, supporting further preclinical and clinical testing of this candidate vaccine.

## Introduction

Since its emergence in December 2019, the Coronavirus Disease 2019 (COVID-19) pandemic has caused more than 90 million infections with more than 2,000,000 deaths worldwide, as of January 15, 2020. The causative agent of COVID-19 was identified to be a novel betacoronavirus (beta-CoV) now known as severe acute respiratory syndrome coronavirus 2 (SARS-CoV-2) (Zhu et al., 2020; Lu et al., 2020). Similar to other human coronaviruses, such as the Middle East respiratory syndrome coronavirus (MERS-CoV) and SARS-CoV, a zoonotic origin of SARS-CoV-2 has been suggested but has not yet been confirmed (Lu et al., 2020; Wu et al., 2020). SARS-CoV-2 can infect individuals from different age groups and causes a wide spectrum of disease manifestations. Most patients with COVID-19 are either asymptomatic or have mild symptoms including fever, myalgia and cough. However, some patients can suffer from moderate to severe life-threatening acute respiratory distress syndrome (ARDS) with possible fatal outcomes (Rodriguez-Morales et al., 2020; Wang et al., 2020; Jin et al., 2020). The high human-to-human transmission of SARS-CoV-2 poses a major challenge toward controlling its spread (Adhikari et al., 2020; Bai et al., 2020; Xu et al., 2020).

Coronaviruses are enveloped viruses with a positive-sense single-stranded ~30 kb RNA genome, which encodes four structural proteins including the surface spike (S) glycoprotein, the envelope (E) protein, the membrane (M), and the nucleocapsid (N). It also encodes 16 nonstructural proteins (nsp1-16) as well as putative and known accessory proteins involved viral replication and pathogenesis (Jin et al., 2020; Fehr and Perlman, 2015; Tan et al., 2005). The viral S protein is capable of inducing a strong immune response. It is comprised of S1 and S2 subunits in which S1 in involved in binding to Angiotensin-converting enzyme 2 (ACE2) receptor on host cells and S2 being involved in viral-host membranes fusion (Yan et al., 2020; Du et al., 2009). Therefore, neutralizing antibodies (nAbs) mainly function through targeting the receptor binding domain (RBD) in the S1 subunit and preventing viral entry in host cells. Indeed, a strong association between the magnitude of anti-S nAbs response and patient survival was recently shown in COVID-19 patients (Sun et al., 2020). Furthermore, the recently approved vaccines for emergency use against SARS-CoV-2 including either mRNA- or adenovirus-based vaccines also target the S protein of the virus. Such work as well as other studies on MERS-CoV and SARS-CoV suggest that viral S protein could represent the main target for the development of vaccines against SARS-CoV-2 (Prompetchara et al., 2020; Fukushi et al., 2018; Fukushi et al., 2006). This is further supported by the isolation and development of several therapeutic human nAbs against the SARS-CoV-2 S protein and their ability to neutralize and block viral entry and/or cell-cell spread at very low concentrations, and sometimes to confer prophylactic and therapeutic protection in animals and humans (Papageorgiou and Mohsin, 2020; Hussen et al., 2020).

The ideal strategy for rapidly controlling existing and potential SARS-CoV-2 outbreaks is to develop a safe and effective vaccine. Several vaccines candidates based on full-length or truncated S protein are being developed and investigated including DNA vaccines, RNA vaccines, replicating or non-replicating viral vectored vaccines, nanoparticle-based vaccine, whole inactivated vaccine (WIV), and S or RBD protein-based subunit vaccines (Zhang et al., 2020; Dong et al., 2020; Krammer., 2020). Many of these vaccines are at the late stages of the clinical trials and/or approved for emergency use in several countries.

Synthetic DNA vaccine is a fast and an easy platform for the production of vaccines compared to other vaccine development technologies due to their simple design, timely production, manufacturing scalability and easy and well-established quality control in addition to their temperature stability (Williams et al., 2013). In addition, DNA vaccines can elicit Th1-biased immune response in contrast to other vaccine platforms such as protein-based subunit vaccines which could induce the undesired Th2-skewed response in the case of coronaviruses (Tseng et al., 2012; Agrawal et al., 2016).

Our previous work showed that DNA vaccines expressing MERS-CoV S1 or full-length S glycoprotein can induce Th1-skewed immune response, nAbs and S-specific T cell response in immunized mice. In this work, we built on our previous experience in developing DNA vaccines for influenza (Hashem et al., 2012) and MERS-CoV (Alamri et al., 2017) to develop a synthetic codon-optimized DNA vaccine as a countermeasure to aid in controlling SARS-CoV-2 spread. Using a plasmid DNA vector suitable for clinical development (pVAX1), here we describe the design and report the results from our preclinical immunogenicity testing of this vaccine candidate, which is denoted as VIU-1005. The design of the VIU-15 vaccine candidate was based on the synthesis of codon-optimized coding sequence for optimal mammalian expression of a consensus full-length S glycoprotein. The successful construction of the vaccine was confirmed, and the expression of the S protein was verified by western blot and immunofluorescence staining. In two mouse models (BALB/c or C57BL/6J mice) we studied the immunogenicity of this candidate vaccine. We show that systemic S-specific IgG antibodies and nAbs were significantly induced in both models post 3 intramuscular needle injections with 100 μg of VIU-1005 vaccine, and such immunization could induce Th1-skewed and long-lasting antibody responses and cellular immunity in mice. We further show that the immunogenicity and induction of Th1-skewed humoral response could be enhanced with the use of needle-free immunization system in which low doses could be sufficient to elicit significant levels of systemic S-specific IgG antibodies and nAbs via intramuscular or intradermal immunization. Our data prove the immunogenicity of our synthetic COVID-19 DNA vaccine and support further preclinical and clinical testing of this candidate vaccine.

## Materials and Methods

### In silico design of codon-optimized synthetic consensus S protein

All available SARS-CoV-2 full-length S protein sequences (399 sequences) by March 10^th^, 2020 were downloaded from GISAID database and dataset was filtered by removing sequences containing ambiguous amino acid codes (BJOUXZ). The final dataset was multiply aligned using CLUSTALW and the Shannon entropy for each amino acid position were determined and the consensus protein sequence was then obtained for the full-length S glycoprotein. The coding sequence for the consensus protein sequence was then codon-optimized for mammalian expression and synthesized by GenScript USA Inc (Piscataway, NJ).

### DNA constructs

The designed full-length codon-optimized consensus coding sequence for SARS-CoV-2 S protein was cloned into the mammalian expression vector pVAX1 under the control of the cytomegalovirus immediate-early promoter and denoted as VIU-1005. The coding sequence was cloned between *NheI* and *KpnI* restriction sites using the T4 DNA ligase (**Figure 1a**). The construct was confirmed by restriction digestion and sequencing. Bulk endotoxin-free preparations of VIU-1005 and the empty control plasmid (control) were prepared for animal studies using GenElute™ HP Select Plasmid Gigaprep Kit (Sigma, Germany).

**Figure 1.**
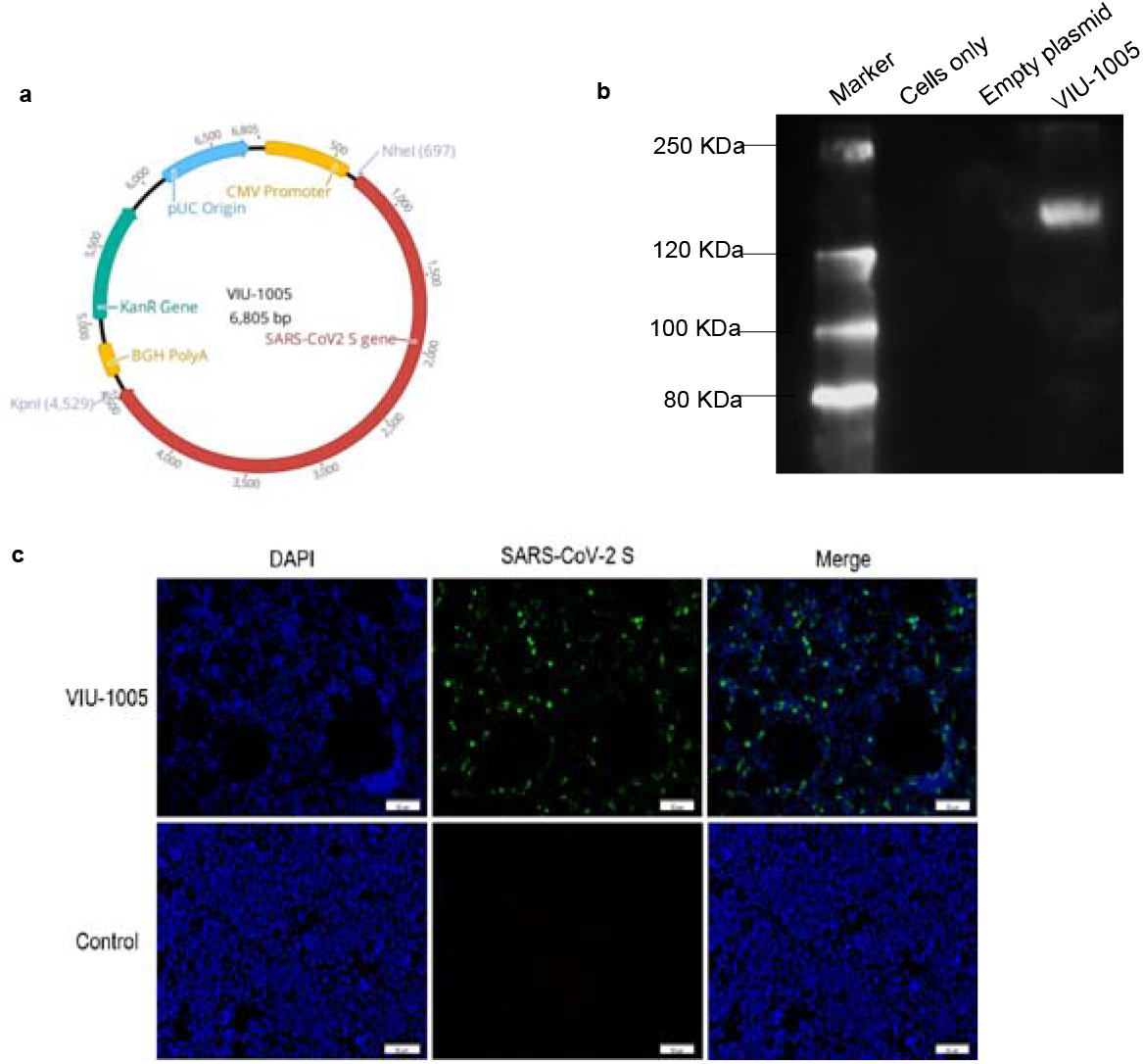
Design and expression of SARS-CoV-2 Spike protein from DNA vaccine. **(a)** Schematic diagram of SARS-CoV-2 DNA vaccine construct (VIU-1005), the SARS-CoV-2 spike gene is indicated by red color. **(b)** Western blot showing the expression of S protein at the expected size from HEK-293 cells transfected with VIU-1005 construct only but not cells only control or cells transfected with empty control plasmid. **(c)** Immunofluorescent staining of cells transfected with VIU-1005 or control plasmid. Transfected cells were stained with anti-SARS-CoV-2 S mouse polyclonal antibodies (green), and nuclei were counterstained with DAPI (blue). Scale bars are 50 μm.

### Cells

Baby Hamster kidney BHK-21/WI-2 cell line (Kerafast, EH1011), African Green monkey kidney-derived Vero E6 cell line (ATCC, CRL-1586) and Human embryonic kidney 293 cells (ATCC, CRL-1573) were cultured in Dulbecco’s modified essential medium (DMEM) containing penicillin (100 U/ml) and streptomycin (100 μg/ml) and supplemented with 5 or 10% fetal bovine serum (FBS) in a 5% CO_2_ environment at 37°C.

### Western blot

Briefly, 70–90% confluent HEK-293 cells in 6-well plates were transiently transfected with 2 μg of VIU-1005 or control plasmid using JetPRIME^®^ Transient Transfection Protocol and Reagents (Polyplus, New York, NY) according to the manufacturer’s instructions. Transfected cells were incubated at 37°C in a 5% CO_2_ incubator for 48 h. Transfected cells were then washed with phosphate-buffered saline (PBS) and lysed with radioimmunoprecipitation assay buffer (RIPA buffer) (Sigma, Germany). The harvested cell lysates were subjected to western blot analysis to verify the expression of S protein using in house mouse anti-S (SARS-CoV-2) polyclonal antibodies at a 1:2000 dilution.

### Immunofluorescence analysis

HEK-293 cells (70% confluent) on a 8-well cell culture slide [growth area/well (cm²): 0.98 and working volume/well (ml): 0.20 - 0.60] were transfected with 0.2 μg of VIU-1005 or control plasmid using JetPRIME^®^ Transient Transfection Protocol and Reagents (Polyplus, New York, NY) according to manufacturer’s instructions, and followed incubated at 37°C in a 5% CO_2_ incubator for 24 h. The media was removed, and the cells were washed with PBS and fixed with 4% formaldehyde at 4°C for 10 min. Cells were then washed twice with PBS and permeabilized with 0.2% PBS-T (Triton 100) at 4°C for 20 min. Fixed cells were washed twice with PBS-T, and the nonspecific binding was blocked with 2% goat serum in PBS-T at room temperature for 30 min and washed twice with PBS-T for 5 min. Cells were then incubated with mouse anti-SARS-CoV-2 S polyclonal antibodies in blocking buffer at 1:2000 dilution at 4°C overnight. After three washes with PBS-T, cells were incubated with Alexa Fluor-488 labeled goat anti-mouse IgG H&L secondary antibody (Abcam, UK) at 1:500 dilution in blocking buffer in the dark at room temperature for 1 h. Cells were washed again for three times with PBS-T, and slides were mounted with VECTASHIELD antifade mounting medium with DAPI counter stain (Invitrogen, Carlsbad, CA). Images were captured using Olympus BX51 Fluorescence Microscope and were analyzed using Image-Pro Plus software.

### Animal Studies

Six to 8-week-old female BALB/c or C57BL/6J mice were obtained from and housed in the animal facility in King Fahd Medical Research Center (KFMRC), King Abdulaziz University (KAU), Jeddah, Saudi Arabia. All animal experiments were conducted in accordance with the guidelines and approval of the Institutional Animal Care and Use Committee (IACUC) at KFMRC and the ethical approval from the bioethical committee at KAU (approval number 04-CEGMR-Bioeth-2020). In one experiment, two groups of BALB/c or C57BL/6J mice (10 per group) were intramuscularly immunized via needle injection with 3 doses of 100 μg of either VIU-1005 or control plasmid at 2-week interval and blood samples were collected for serological testing every 2 weeks starting from day 0 (pre-bleed) until week 8. In another experiment, three groups of BALB/c mice (5 per group) were immunized intramuscularly, intradermally or subcutaneously via needle injection with 3 doses of 100 μg of either VIU-1005 or control plasmid at 2-week interval and blood samples were collected at different time points until week 25 post primary immunization. In a third experiment, BALB/c mice (4-5 per group) were intramuscularly or intradermally immunized with 3 doses of 25 μg, 50 μg or 100 μg of VIU-1005 plasmid at 2-week interval using either conventional needle injection or customized needle-free Tropis system (PharmaJet, Golden, CO) and blood samples were collected every week until week 8 post primary immunization. Spleens were collected from some animals for flow cytometry analysis as outlined below.

### Indirect ELISA

The end-point titers or optical density (OD) readings at 1:100 dilution of total anti-S1 IgG or its isotypes (IgG1, IgG2a and IgG2b) from immunized mice were determined by enzyme-linked immunosorbent assay (ELISA) as described previously (Alamri et al., 2017). Briefly, 96-well EU Immulon 2 HB plates (Thermo Scientific) were coated overnight at 4°C with the SARS-CoV-2 S1 subunit (amino acids 1–685) (Sino Biological, China) at 1 μg/ml in PBS (50 ul/well). Then, the plates were washed three times with washing buffer (PBS containing 0.1% Tween-20 (PBS-T)). This was followed by blocking with 200 μl/well of blocking buffer (5% skim milk in PBS-T) for 1 h at room temperature. Plates were washed three times and incubated with a 2-fold serial dilution of mouse sera (100 μl/well) starting from 1:100 dilution in blocking buffer and incubated for 1 h at 37°C. Some samples collected at different time points were only tested at 1:100 dilution. After three washes, peroxidase-conjugated rabbit anti-mouse IgG secondary antibodies as well as anti-IgG1, IgG2a or IgG2b antibodies (Jackson Immunoresearch Laboratories, West Grove, PA) were added at dilutions recommended by the manufacturer and incubated for 1 h at 37°C as 100 μl/well. Excess secondary antibodies were removed by three washes and color was developed by adding 3,3’,5,5’-Tetramethylbenzidine (TMB) substrate (KPL, Gaithersburg, MD) for 15 min. Finally, reactions were stopped with 0.16 M sulfuric acid and absorbance was read spectrophotometrically at 450 nm using the ELx808™ Absorbance Microplate Reader (BioTek, Winooski, VT). End-point titers were determined and expressed as the reciprocals of the highest dilution with OD reading above the cut-off value defined as the mean of the control group plus three standard deviations (SD).

### SARS-CoV-2 pseudovirus neutralization assay

Pseudovirus microneutralization assay was performed as previously described (Almahboub et al., 2020). Briefly, rVSV-ΔG/SARS-2-S*-luciferase pseudovirus was generated by transfecting BHK21/WI-2 cells with pcDNA expressing codon-optimized full-length SARS-CoV-2 S protein (GenBank accession number: MN908947) using Lipofectamine^™^ 2000 transfection reagent (Invitrogen, Carlsbad, CA). Transfected cells were then infected with rVSV-ΔG/G*-luciferase 24 h later, and the supernatant containing the generated rVSV-ΔG/SARS-2-S*-luciferase pseudovirus was collected 24 h post-infection. The collected virus was titrated by measuring luciferase activity from serially diluted supernatant on Vero E6 cells and the titer was expressed as a relative luciferase unit (RLU). Neutralization assay was then conducted by incubating two-fold serial dilutions of heat-inactivated mouse sera from vaccinated and control groups starting from 1:20 dilution (in duplicate) with DMEM-5%FBS containing 5×10^4^ RLU rVSV-ΔG/SARS-2-S*-luciferase pseudovirus for 1 h at 37 °C in a 5% CO_2_ incubator. Pseudovirus–serum mixtures were transferred onto confluent Vero E6 cell monolayers in white 96-well plates and incubated for 24 h at 37°C in a 5% CO_2_ incubator. After 24 h, cells were lysed, and luciferase activity was measured using Luciferase Assay System (Promega) according to the manufacturer’s instructions, and the luminescence was measured using BioTek Synergy 2 microplate reader (BioTek, Winooski, VT). Cell only control (CC) and virus control (VC) were included with each assay run. The median inhibitory concentration (IC_50_) of neutralizing antibodies (nAbs) was determined using four-parameter logistic (4PL) curve in GraphPad Prism V8 software (GraphPad Co.) and calculated as the reciprocal of the serum dilution at which RLU was reduced by 50% compared with the virus control wells after subtraction of the background RLUs in the control groups with cells only.

### Flow cytometry

Memory CD8^+^ and CD4^+^ T cells interferon (IFN-γ), tumor necrosis factor–α (TNF-α) and interleukin-2 (IL-2) IL-2 responses were evaluated at 4 weeks after last immunizations from immunize animals. Single-cell suspensions of splenocytes were prepared from individual immunized and control BALB/c mice. In brief, spleens from mice were collected in 3 ml of RPMI 1640 (Invitrogen, Carlsbad, CA) supplemented with 5% heat inactivated FBS and smashed between frosted ends of two glass slides. Processed splenocytes were then filtered through 70-μm nylon filters and centrifuged at 430 x g for 5 min at room temperature. Red blood cells were then lysed by adding 3 ml of ammonium-chloride-potassium (ACK) lysis buffer (Invitrogen, Carlsbad, CA) for 4 min at room temperature, and equal volume of PBS was then added for neutralization. Cells were centrifuged again at 430 x g for 5 min at room temperature and cell pellets were resuspended in RPMI 1640 at a concentration of 1 × 10^7^ cells/ml. One million cells per well were added to a U-bottom 96-well plate and were stimulated with 5 μg/ml of pools of overlapping SARS-CoV-2 S protein peptides (GenScript USA Inc, Piscataway, NJ). The stimulation was performed by incubation for 6 h at 37°C and 5% CO_2_ in the presence of Protein Transport Inhibitor Cocktail (brefeldin A) (BD Biosciences, San Jose, CA) at a final concentration of 1:1000. Splenocytes stimulated in the presence of phorbol 12- myristate 13-acetate (PMA), ionomycin, and brefeldin A were used as a positive control, and unstimulated splenocytes in RPMI 1640 medium were used as a negative control. Cells were then washed in FACS buffer (PBS with 2% heat inactivated FBS) and stained with LIVE/DEAD Fixable Violet Dead Cell Stain Kit (Invitrogen, Carlsbad, CA) for 30 min at room temperature. After washing with PBS, PB-conjugated anti-mouse CD8, PB-conjugated anti-mouse CD4, APC-conjugated anti-mouse CD44 antibody and Pe-Cy7-conjugated anti-mouse CD62L antibodies (BioLegend, UK) were used for surface markers staining. The cells were then washed with FACS buffer and fixed and permeabilized using Cytofix/Cytoperm Solution (BD Biosciences, San Jose, CA) according to the manufacturer’s protocol. For intracellular staining, cells were labeled with APC-Cy7-conjugated anti–mouse IFN-γ (clone XMG1.2), PE-conjugated anti-mouse TNF-α (clone MP6-XT22) and Pe-Cy7–conjugated anti-mouse IL-2 (clone JES6-5H4) antibodies (BioLegend, UK) for 20 min at 4°C. Cells were then washed twice with permeabilization buffer and once with FACS buffer. All data were collected using a 15 laser Aria flow cytometer (BD Biosciences, San Jose, CA) and analyzed using FlowJo v10 software (Tree Star).

### Statistical analysis

Statistical analyses and graphical presentations were conducted using GraphPad Prism version 8.0 software (Graph-Pad Software, Inc., CA, USA). Statistical analysis was conducted using one-way analysis of variance with Bonferroni post-hoc test to adjust for multiple comparisons between groups, or Mann–Whitney test. All values are depicted as mean ± SD and statistical significance is reported as *, P≤0.05, **, P≤0.01, ***, *p* ≤ 0.001, and ****, *p* ≤ 0.0001.

## Results

### In vitro protein expression from the synthetic DNA vaccine

Prior to animal experiments, spike protein expression from the VIU-1005 construct (**Figure 1a**) was confirmed *in vitro* in HEK-293 cells. Western blot analysis confirmed that the recombinant construct was able to express the spike protein indicated by the band observed at expected molecular weight, but not in lysates from cells transfected with the control plasmid as shown in **Figure 1b**. Immunofluorescence analysis was also performed to visualize the expression and localization of SARS-CoV-2 Spike protein in transfected HEK-293 cells. As shown in **Figure 1c**, a strong S protein expression in the cytoplasm was only detected in cells transfected with VIU-1005 using mouse anti-SARS-CoV-2 S polyclonal antibodies but not in cells transfected with the control plasmid.

### Intramuscular immunization with VIU-1005 vaccine elicits significant Th1-skewed humoral immune response in two mouse models

Next, we investigated the immunogenicity induced by the generated synthetic DNA vaccine candidate (VIU-1005) in BALB/c and C57BL/6J mice (**Figures 2a and 3a**). The two mouse models were used to evaluate the type and nature of the immune response as BALB/c and C57BL/6J mice are phenotypically represent Th2 and Th1 skewed models, respectively. Mice intramuscularly immunized with three doses of the vaccine induced significant levels of S1-specific IgG. Specifically, while we observed that immunization with two doses elicited significant levels of S1-specific IgG in both BALB/c and C57BL/6J mice (i.e. on week 4), samples collected 4 weeks post-third immunization (i.e. on week 8) showed higher and more significant levels compared to control groups immunized with the control plasmid (**Figures 2b and 3b**). Interestingly, immunization with VIU-1005 significantly induced higher levels of S1-specific IgG2a and IgG2b isotypes compared to IgG1 in both animal models (**Figures 2c and 3c**). As expected, the bias towards Th1 response was more pronounced in C57BL/6J mice since they are Th1-prone animals compared to BALB/c mice as shown by the high IgG2a/IgG1 or IgG2b/IgG1 ratios (**Figures 2d and 3d**). Nonetheless, BALB/c mice also showed Th1-skewed response, confirming that VIU-1005 could elicit elevated titers of S1-specific IgG2a and IgG2b antibodies which is indicative of a skewed Th1 immune response despite the genetic and phenotypic background of the animal model.

**Figure 2.**
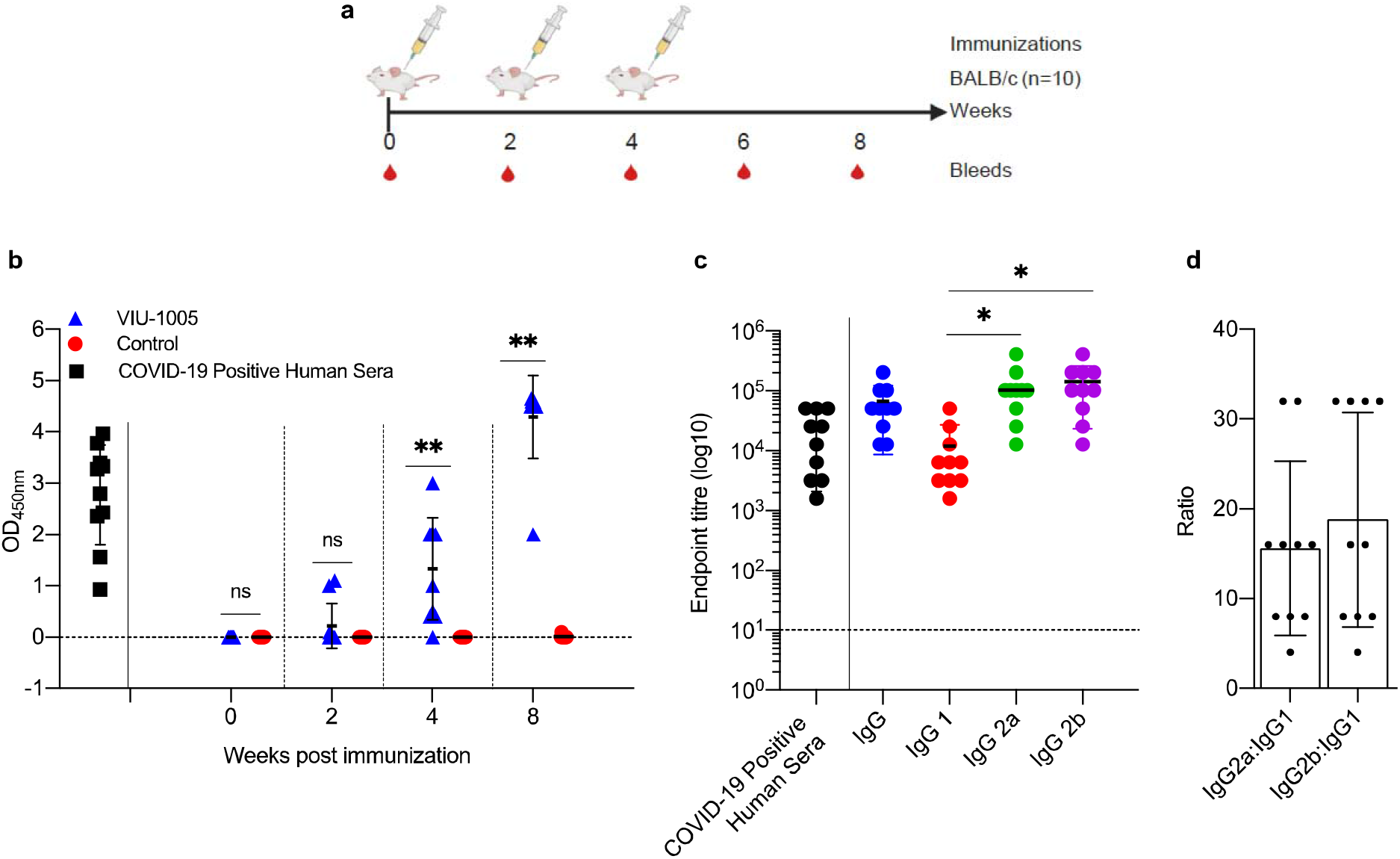
Antibody response against SARS-CoV-2 S in BALB/c mice. (**a**) Mice were intramuscularly immunized with 3 doses of 100 μg at 2-week interval using either VIU-1005 or control plasmid. The timeline shows the bleeding and immunization regimen. (**b**) OD values of S1-specific binding total IgG at 1:100 dilution from each mouse were determined by ELISA at 2, 4 and 8 weeks post first immunization on day 0. IgG response from recovered COVID-19 human patients is also shown to compare the data with the immunized mice sera. (**c**) End-point titers of S1-specific total IgG, IgG1, IgG2a and IgG2b were determined by ELISA in samples collected on week 8 from immunized mice. End-point titers of total IgG from recovered COVID-19 human patients is also shown to compare the data with the immunized mice sera. (**d**) IgG2a:IgG1 and IgG2b:IgG1 ratios were calculated in samples collected from immunized mice on week 8. Data are shown as mean ± SD for each group from one experiment (n = 10). Statistical significance was determined by Mann–Whitney test in (**c**) and one-way analysis of variance with Bonferroni post-hoc test in (**d**).

**Figure 3.**
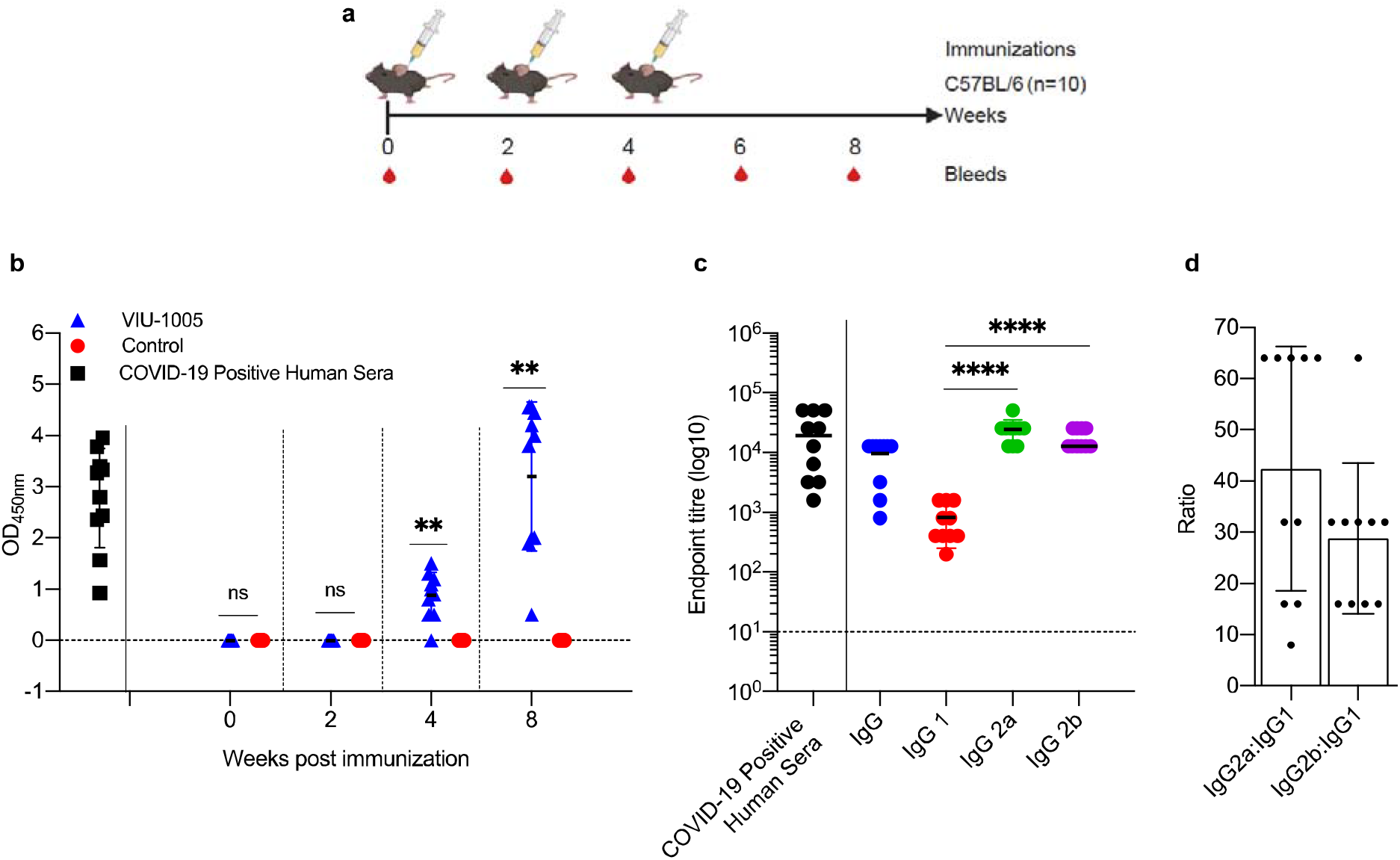
Antibody response against SARS-CoV-2 S1 in C57BL/6J mice. (**a**) Mice were intramuscularly immunized with 3 doses of 100 μg at 2-week interval using either VIU-1005 or control plasmid. The timeline shows the bleeding and immunization regimen. (**b**) OD values of S1-specific binding total IgG at 1:100 dilution from each mouse were determined by ELISA at 2, 4 and 8 weeks post first immunization on day 0. IgG response from recovered COVID-19 human patients is also shown to compare the data with the immunized mice sera. (**c**) End-point titers of S1-specific total IgG, IgG1, IgG2a and IgG2b were determined by ELISA in samples collected on week 8 from immunized mice. End-point titers of total IgG from recovered COVID-19 human patients is also shown to compare the data with the immunized mice sera. (**d**) IgG2a:IgG1 and IgG2b:IgG1 ratios were calculated in samples collected from immunized mice on week 8. Data are shown as mean ± SD for each group from one experiment (n = 10). Statistical significance was determined by Mann–Whitney test in (**c**) and one-way analysis of variance with Bonferroni post-hoc test in (**d**).

### Intramuscular immunization with VIU-1005 vaccine induces neutralizing antibodies in both BALB/c and C57BL/6J mice

To further investigate the effector function of the elicited antibodies generated by the developed vaccine, sera from immunized and control mice were tested in pseudovirus microneutralization assay to evaluate nAbs. As shown in **Figure 4**, sera collected on week 8 from VIU-1005 immunized group induced significant levels of nAbs compared to control group with mean IC_50_ titers of 1×10^3^ in both BALB/c (**Figures 4a and 4b**) and C57BL/6J (**Figures 4c and 4d**) mice. As expected, no neutralizing activity was observed from mice immunized with control plasmid.

**Figure 4.**
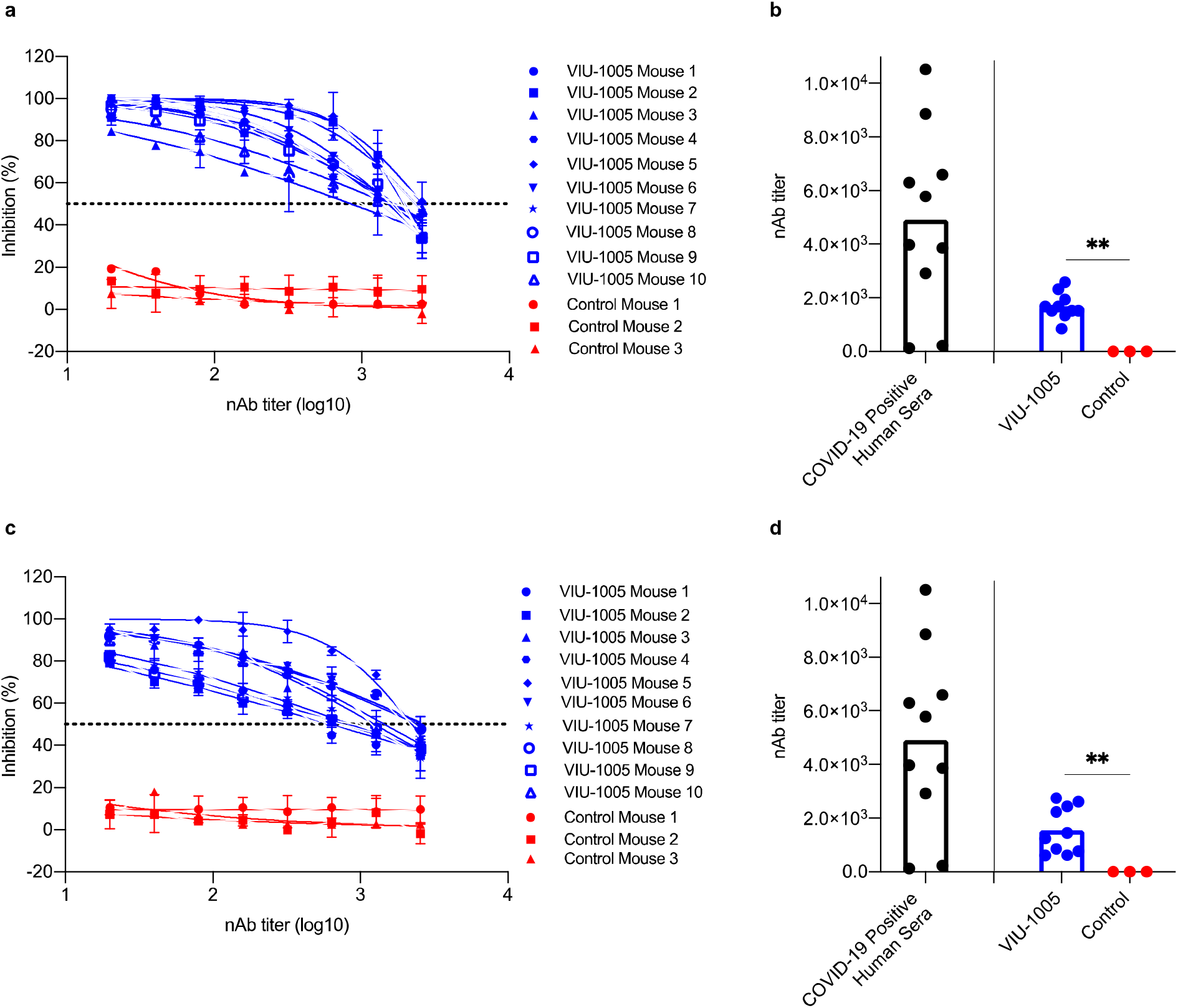
Neutralizing antibody response after intramuscular immunization in BALB/c and C57BL/6J mice. (**a and b**) BALB/c or (**c and d**) C57BL/6J mice were intramuscularly immunized with 3 doses of 100 μg at 2-week interval using either VIU-1005 (n = 10) or control plasmid (n = 3). Serum samples were collected at week 8 post first immunization and serially diluted and tested in duplicate for their neutralizing activity against rVSV-ΔG/SARS-2-S*-luciferase pseudovirus as described in materials and methods. (**a and c**) Neutralizing activity from serum samples collected from each mouse, and (**b and d**) nAb titers (IC_50_). nAbs from recovered COVID-19 human patients were used as a control (**b and d**) for the assay and to compare the data with the immunized mice sera. Data are shown as mean ± SD in (**a and c**) and bar represent the mean in (**b and d**) from one experiment. Statistical significance was determined by Mann–Whitney test in (**b and d**).

### Intramuscular immunization with VIU-1005 vaccine elicits long lasting humoral immune response in both BALB/c and C57BL/6J mice

To determine if route of immunization can affect immunogenicity of our vaccine candidate and to measure the longevity of the generated antibody response in mice, we immunized BALB/c mice with different immunization routes and compared the immunogenicity of VIU-1005 over extended period of time. Interestingly, intradermal and subcutaneous immunization of BALB/c mice with three doses of VIU-1005 failed to induce any significant IgG levels (**Figures 5c and 5d**). On the other hand, such immunization via intramuscular route elicited long-lasting S1-specific IgG that lasted until week 25 post-primary immunization (**Figure 5a**). Taken together, these data suggest that intramuscular immunization with this synthetic codon-optimized DNA vaccine could induce long-lasting antibody response in mice.

**Figure 5.**
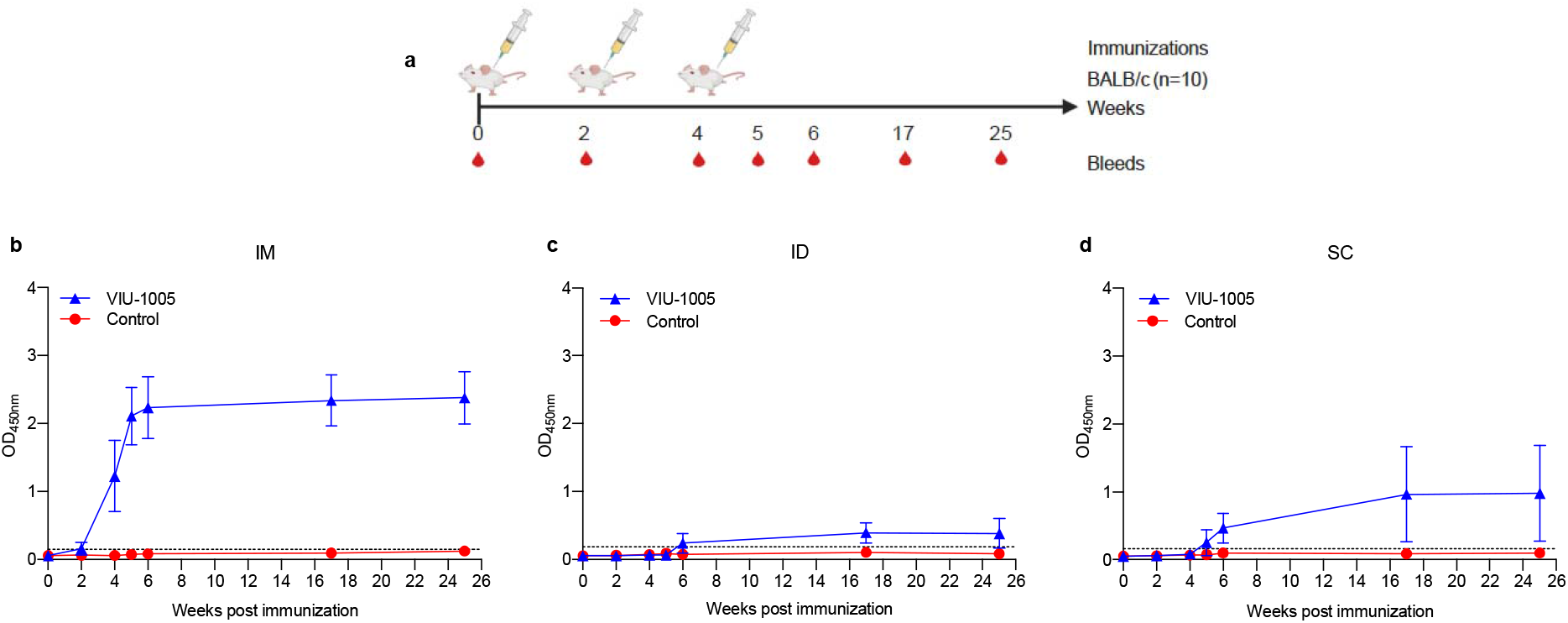
Long-term IgG response in mice immunized with different routes. (**a**) BALB/c mice were immunized via different routes using 3 doses (100 μg each) at 2-week interval using either VIU-1005 or control plasmid. The timeline shows the bleeding and immunization regimen. OD values of S1-specific binding total IgG at 1:100 dilution from each mouse immunized via (**b**) intramuscular, (**c**) intradermal, or (**d**) subcutaneous routes were determined by ELISA at 2, 4, 5, 6, 17 and 25 weeks post first immunization on day 0. Data are shown as mean ± SD for each group from one experiment (n = 5).

### Needle-free intramuscular immunization enhances the immunogenicity of VIU-1005 vaccine in mice

To further improve the immunogenicity of the naked synthetic DNA vaccine and minimize the number and size of doses, we investigated the use of needle-free Tropis system with our vaccine in BALB/c mice (**Figure 6)**. Immunization with only two or three doses of as low as 25 μg of the vaccine via intramuscular route was able to elicit significantly high levels of S1-specific total IgG. Specifically, two or three doses of VIU-1005 were able to induce high levels of S1-specific total IgG in a dose-dependent fashion (**Figure 6b**), in which two doses of 50 μg and 100 μg administered by the needle-free system elicited S1-specific total IgG levels that are equivalent or higher than that induced by three doses of 100 μg by needle injection shown in **Figure 2b**. High levels of endpoint titers of S1-specific total IgG from week 8 sera of mice intramuscularly immunized with three doses of 25 μg, 50 μg and 100 μg of VIU-1005 were also observed (**Figure 6c**). As expected, no S1-specific IgG antibody response was observed from mice immunized with control plasmid. While we observed a dose-dependent immune response in immunize animals, no significant differences were observed for S1-specific total IgG among the sera of mice immunized with VIU-1005 at 25 μg, 50 μg and 100 μg from week 4 onward. We further investigated the neutralizing activity of the sera from BALB/c mice immunized with the different doses of VIU-1005 vaccine using needle-free Tropis system. As shown in **Figure 6d**, results demonstrated that all three tested doses were able to induce potent titers of nAbs in a dose-dependent manner against SARS-CoV-2 pseudovirus in Vero cells. Immunization with three doses of as low as 25 μg of the vaccine via intramuscular route administered by the needle-free system was sufficient to induce high levels of nAbs, reaching 1×10^3^, which are equivalent to that induced by three doses of 100 μg by needle injection (**Figure 4b**). These results suggest that 25 or 50 μg of VIU-1005 is sufficient to induce high titers of S1-specific antibodies and nAbs in mice using the needle-free system.

**Figure 6.**
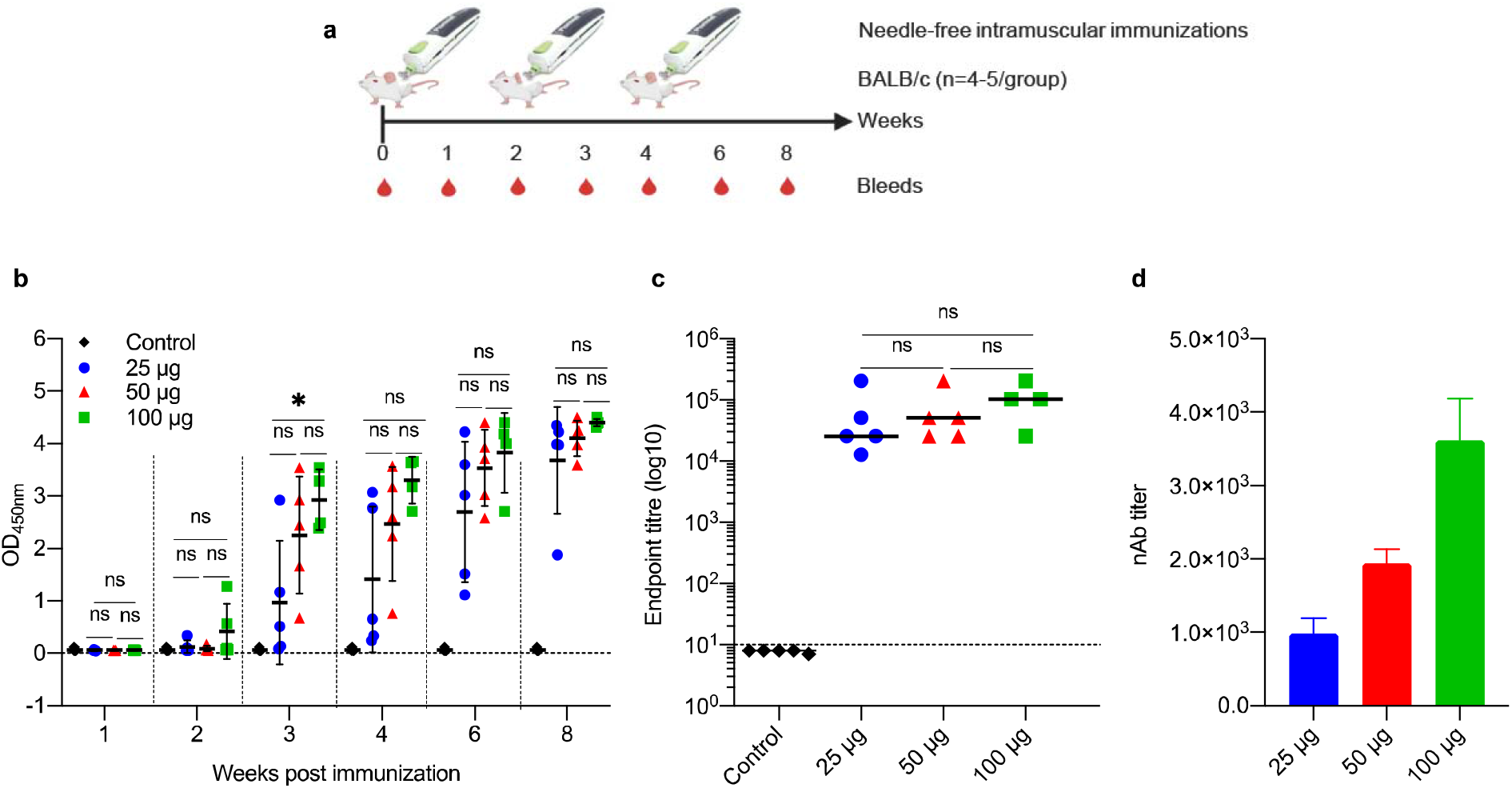
Antibody response in BALB/c mice intramuscularly immunized with VIU-1005 using needle-free Tropis system. (**a**) Timeline of bleeding and immunization regimen. BALB/c mice were immunized intramuscularly with 3 doses of 25 μg, 50 μg or 100 μg at 2-week interval using VIU-1005 vaccine or control plasmid. (**b**) OD values of S1-specific binding total IgG at 1:100 dilution from each mouse were determined by ELISA at 1, 2, 3, 4, 6 and 8 weeks post first immunization on day 0. (**c**) End-point titer of S1-specific total IgG was determined by ELISA in samples collected from immunized mice at week 8. (**d**) nAb titers (IC_50_) from mice immunized with doses of 25 μg, 50 μg or 100 μg of VIU-1005 plasmid using needle-free system were determined at week 8. Data are shown as mean ± SD from one experiment (n = 4-5).

To further confirm the superiority of the needle-free system compared to the conventional needle injection, we compared S1-specific total IgG responses in sera collected from BALB/c mice immunized with 25 μg (**Figure 7a**), 50 μg (**Figure 7b**) and 100 μg (**Figure 7c**) using both methods of immunization. As shown in **Figure 7**, needle-free administration of 50 μg and 100 μg not only induced significant high levels of S1-specific total IgG compared to needle injection but we also observed that all mice immunized with the needle-free system elicited S1-specific IgG compared to few from the group immunized using needle injection. Furthermore, needle-free immunization with VIU-1005 significantly induced higher levels of S1-specific IgG1, IgG2a and IgG2b isotypes compared to the conventional needle injection in BALB/c mice, with an elevated titer of IgG2a and IgG2b antibodies suggesting a bias toward Th1 response at the three tested doses (**Figure 8**). Together, these results showed that needle-free immunization not only enhanced S1-specific IgG levels but also boosted Th1 responses, as demonstrated by markedly elevated titers of S1-specific IgG2a and IgG2b antibodies.

**Figure 7.**
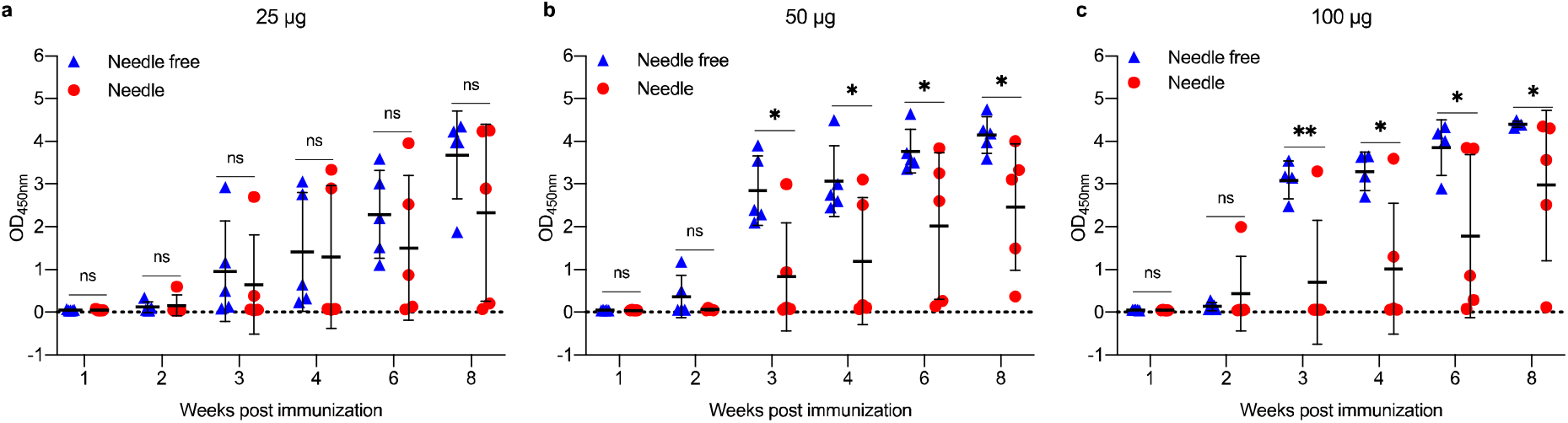
Antibody response against SARS-CoV-2 S1 in BALB/c mice intramuscularly immunized using either needle injection or needle-free Tropis system. BALB/c mice were immunized with 3 doses of (**a**) 25 μg, (**b**) 50 μg or (**c**) 100 μg of VIU-1005 plasmid at 2-week interval using either needle injection or needle-free Tropis system. Binding of total S1-specific IgG at 1:100 dilution from each mouse was determined by ELISA at 1, 2, 3, 4, 6 and 8 weeks post first immunization on day 0. Data are shown as mean ± SD from 4-5 mice from experiment in Figure 6. Statistical significance was determined by Mann–Whitney test.

**Figure 8.**
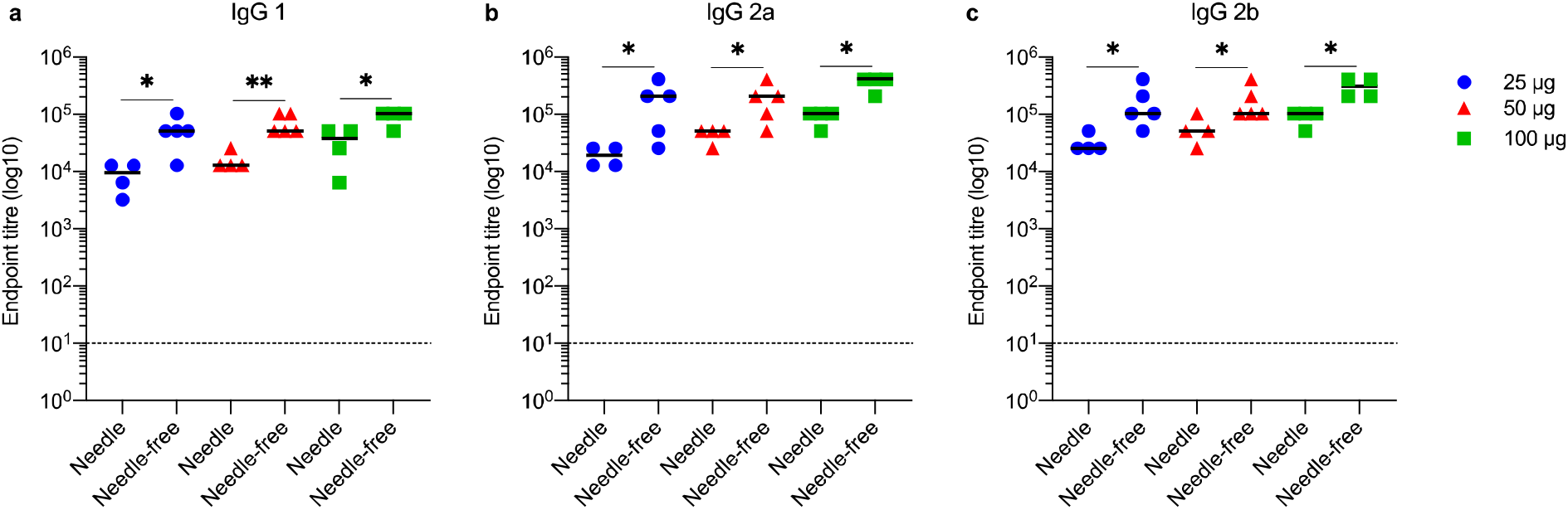
Endpoint antibody titers of IgG isotypes against SARS-CoV-2 S1 in BALB/c mice immunized using either needle injection or needle-free Tropis system. BALB/c mice were immunized intramuscularly with 3 doses of 25 μg, 50 μg or 100 μg of VIU-1005 plasmid at 2-week interval using either needle injection or needle-free Tropis system. End-point titer of S1-specific (**a**) IgG1, (**b**) IgG2a and (**c**) IgG2b were determined by ELISA in samples collected on week 8 from immunized mice. Data are shown as mean from 4-5 mice from experiment in Figures 6 and 7. Statistical significance was determined by Mann–Whitney test.

### Needle-free intradermal immunization with VIU-1005 vaccine induces robust S1-specific humoral response in mice

Interestingly, while intradermal and subcutaneous needle injection using three doses of 100 μg failed to induce high or significant levels of specific IgG (**Figures 5c and 5d**), two or three doses of VIU-1005 administered intradermally via the needle-free system induced significantly high levels of S1-specific IgG in a dose-dependent fashion (**Figure 9b**). Importantly, the levels of S1-specific IgG were equivalent to those generated by three doses administered by intramuscular needle injection. When endpoint IgG titers from the week 8 sera of mice immunized with the three doses of 25 μg, 50 μg and 100 μg were compared, we observed an induction of high levels of IgG antibody response in mice immunized intradermally with 25 μg of VIU-1005 to levels similar to those observed in mice immunized with higher doses with no significant differences (**Figure 9c**). Importantly, such immunization also elicited high titers of nAbs (**Figure 9d**). These data clearly suggest that intradermal needle-free immunization even with low dose could elicit potent humoral immunity in mice, further confirming the superiority of the needle-free system compared to the conventional needle injection.

**Figure 9.**
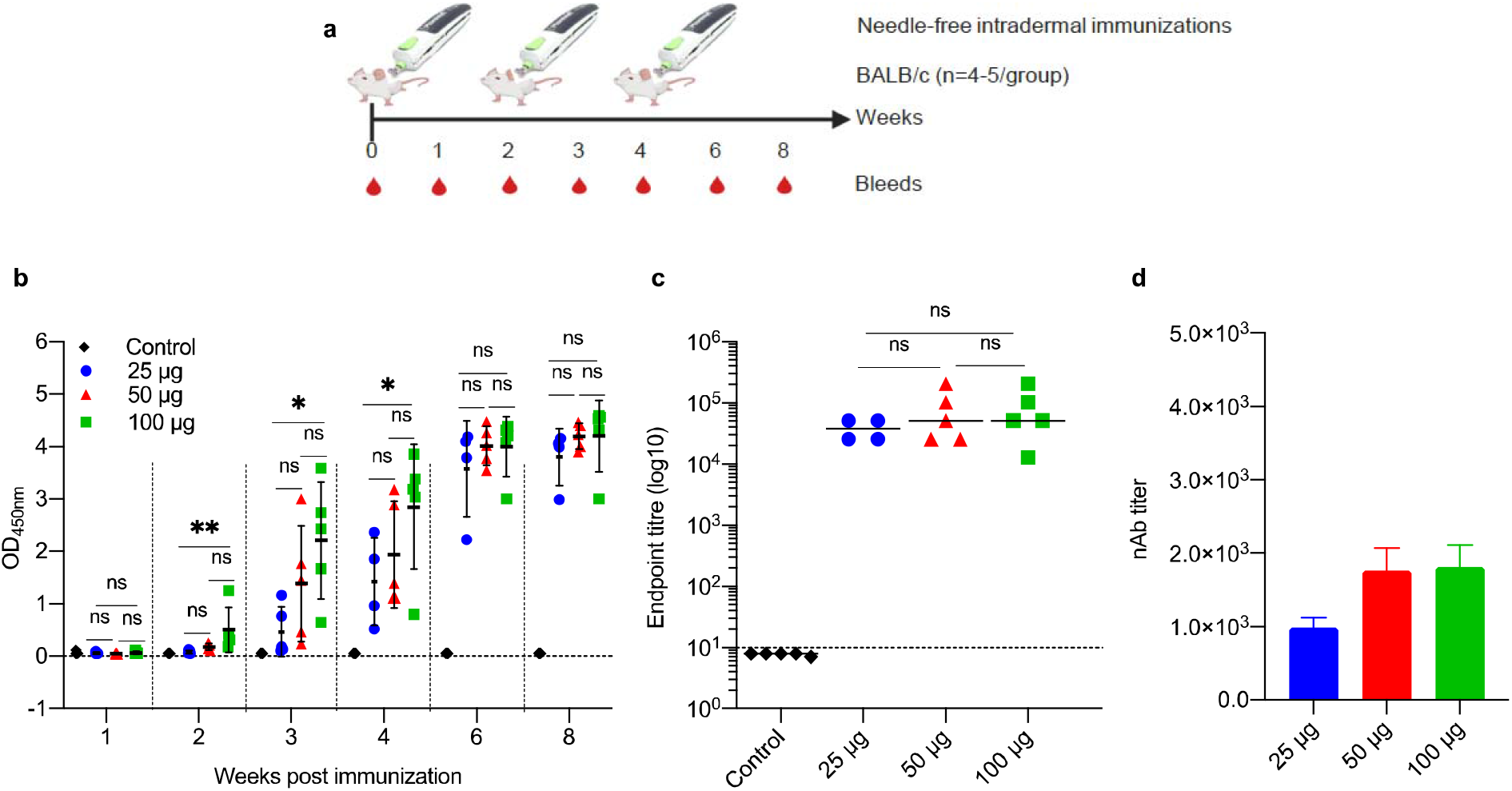
Antibody response against SARS-CoV-2 S1 in BALB/c mice intradermally immunized with VIU-1005 using needle-free Tropis system. (**a**) Timeline of bleeding and immunization regimen. BALB/c mice were immunized intradermally with 3 doses of 25 μg, 50 μg or 100 μg at 2-week interval using VIU-1005 vaccine or control plasmid. (**b**) OD values of S1-specific binding total IgG at 1:100 dilution from each mouse were determined by ELISA at 1, 2, 3, 4, 6 and 8 weeks post first immunization on day 0. (**c**) End-point titer of S1-specific total IgG was determined by ELISA in samples collected from immunized mice at week 8. (**d**) nAb titers (IC_50_) from mice immunized with doses of 25 μg, 50 μg or 100 μg of VIU-1005 plasmid using needle-free system were determined at week 8. Data are shown as mean ± SD from one experiment (n = 4-5).

### Induction of strong S-specific T cell responses in mice immunized with VIU-1005 vaccine

Having observed the Th1-skewed response in VIU-1005 immunized mice, we further tried to evaluate S-specific memory CD8^+^ and CD4^+^ T cells response. To this end, splenocytes isolated from BALB/c mice immunized with 100 μg of VIU-1005 using needle injection were re-stimulated *ex vivo* with four SARS-CoV-2 peptide pools covering SARS-CoV-2 S protein. Flow cytometric analysis showed high levels of IFN-γ, TNF-α and IL-2 memory CD8^+^ and CD4^+^ T cells in VIU-1005 immunized animals only compared to the group immunized with the control plasmid (**Figure 10**), confirming that this vaccine is capable of inducing strong humoral and cellular immune responses in mice.

**Figure 10.**
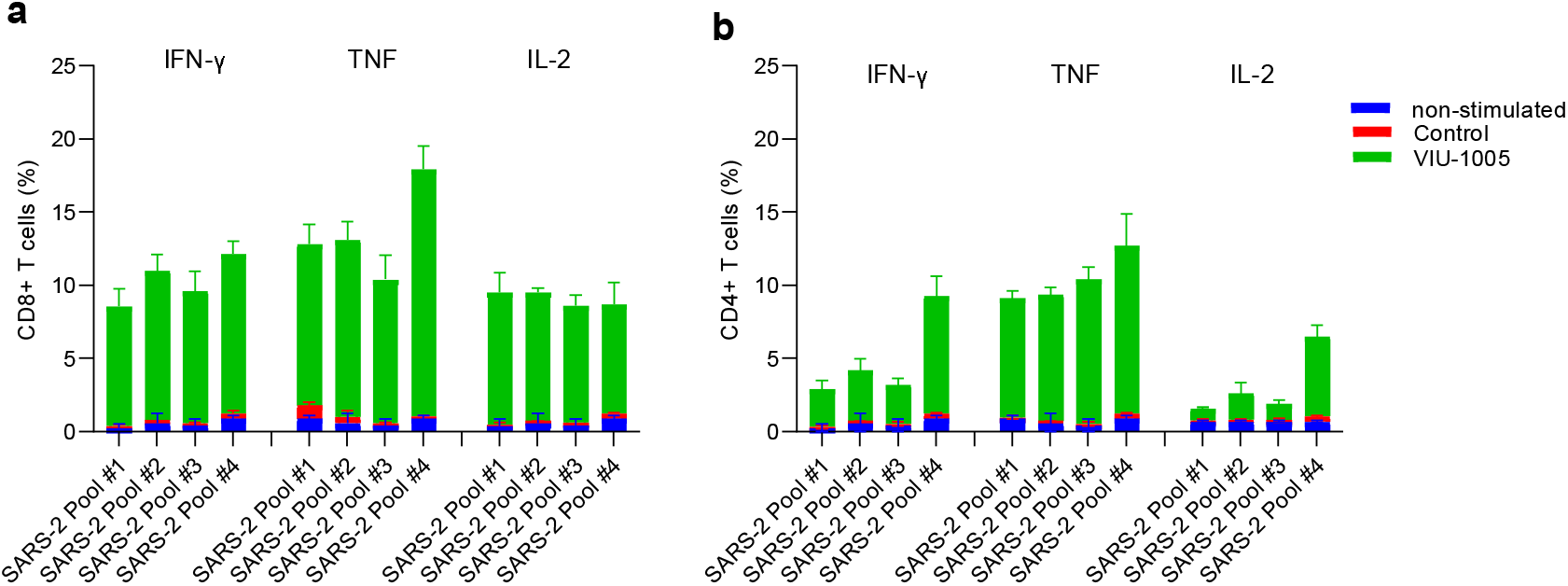
Cellular immune response against SARS-CoV-2 S protein in BALB/c mice. Intramuscularly immunized BALB/c mice with 100 μg of VIU-1005 or control plasmid using needle injection were sacrificed at 4 weeks after last immunization and splenocytes (n = 3) were isolated and re-stimulated ex vivo with synthetic peptide pools covering SARS-CoV-2 S protein. Histograms displaying, IFN-γ expression, TNF-α expression and IL-2 expression on non-stimulated and stimulated (**a**) Memory CD8^+^ cells and (**b**) memory CD4^+^ T cells from immunized BALB/c mice. Data are shown as mean ± SD for each group from one experiment out of two independent experiments (n=3).

## Discussion

There is an urgent need to develop a safe and protective vaccine against SARS-CoV-2 to control the COVID-19 pandemic. Synthetic DNA vaccines represent a promising vaccine platform to use in response to outbreaks, such as COVID-19. They could be quickly designed and synthesized based on viral sequences. Their manufacturing is easy and scalable unlike other platforms such as viral vector or virus based vaccines (Silveira et al., 2017; Xu et al., 2014). They are very stable at different storage conditions which can be extremely practical for use in rural regions (Ingolotti et al., 2010). In addition, the use of consensus S could induce broad immune response by using conserved sequences which should cover possible variation in viral sequences. Therefore, here we analyzed all available SARS-CoV-2 S sequences from GISAID database until March 10^th^, 2020. Such sequence could offer protection against broad range of SARS-CoV-2 strains, possibly including emerging strains.

Based on our previous work in developing a vaccine against MERS-CoV as well as other vaccines developed for other coronaviruses, we selected the SARS-CoV-2 S protein as a target because it is a major protein on the surface of the virus and a main target for nAbs. In vivo testing in mice showed intramuscular immunization with three doses of VIU-1005 via needle injection induced significant and long-lasting levels of Th1-skewed immune response S1-specific IgG in BALB/c (Th2-dominant mouse strain) and C57BL/6J (Th1-dominant mouse strain) mice as well as significant levels of nAbs compared to control group with mean IC_50_ titers of 1×10^3^ in both models. Importantly, needle-free immunization with doses of as low as 25 μg of the VIU-1005 via either intramuscular or intradermal routes was able to elicit high levels of S1-specific IgG in a dose-dependent fashion in BALB/c mice. Two or three doses of 50 μg and 100 μg administered by the needle-free system elicited IgG and nAbs levels that are equivalent or higher than that induced by three doses of 100 μg by needle injection in BALB/c mice. Interestingly, using needle-free system enhanced the immunogenicity of the VIU-1005 vaccine and induced significant levels of S1-specific IgG even at 50 μg and 100 μg when given intradermally albeit the inability of the vaccine to induce any levels of antibodies when administered intradermally via needle injection. Importantly, our findings show that this vaccine could induce long-lasting S1-specific antibodies in mice that were detected for at least 25 weeks after primary immunization.

The use of the needle-free system can not only lead to induction of higher immune response but also has other advantages including lower vaccine volume and higher antigen expression. Clinically, needle-free system would be associated with less pain and stress compared to conventional needle system. As pian at the site of injection was one of the most common adverts effects of the currently approved SARS-CoV-2 vaccine, needle-free system represents an ideal alternative to administer vaccines.

Furthermore, this vaccine showed induction of cellular immunity and increase in cytokine production from both memory CD4^+^ and CD8^+^ T cell compartments, including IFN-γ, TNF-α and IL-2 along with significantly elevated titers of IgG2a and IgG2b, suggesting successful induction of Th1-skewed immune responses as recently reported (Smith et al., 2020). The fact that this vaccine induced Th1-biased immune response in both BALB/c and C57BL/6J mice may alleviate the concerns associated with vaccine-induced immunopathology that has been raised for SARS and MERS vaccine candidates. Such immunopathology is characterized by Th2-skewed immune response and eosinophilia and was reported for different vaccines developed for MERS-CoV and SARS-CoV after viral challenge (Weingartl et al., 2004; Yang et al., 2005; Czub et al., 2005; Deming et al., 2006; Yasui et al., 2008; Bolles et al., 2011; Tseng et al., 2012; Jaume et al., 2012; Iwata-Yoshikawa et al., 2014; Honda-Okubo et al., 2015; Agrawal et al., 2016; Hashem et al., 2019).

Taken together, our study confirms the immunogenicity of VIU-1005 vaccine candidate expressing full-length consensus SARS-CoV-2 S glycoprotein in eliciting strong Th1-skewed and long-lasting protective humoral and cellular immunity, and suggest that it could represent a promising protective candidate vaccine against SARS-CoV-2.

## Author Contributions

A.A., M.H. and A.M.H. designed and conceptualized the work. S.S.A., K.A.A., R.Y.A., A.A., S.A., M.A.A., T.S.A., W.A., M.Z.E., R.A. and A.M.H. performed and optimized experiments and analyzed data. S.S.A., K.A.A., A.A., and A.M.H. drafted the manuscript. All authors have reviewed and edited the manuscript and agreed to the published version of the manuscript.

## Acknowledgments

We would like to thank King Fahd Medical Research Center (KFMRC) and King Abdulaziz University (KAU) for their continuous support.

## Funding

This work was supported by King Abdulaziz City for Science and Technology (KACST), Riyadh, Saudi Arabia, through research grant program (number 09-1), which is a part of the Targeted Research Program (TRP).

## Conflict of interest

AMH, MAA, TSA, KAA, SSA and AA are inventors on a US patent application related to this work. AMH and AA work as scientific consultants in SaudiVax ltd and receive fees for consulting. SA and MH are employees of SaudiVax ltd and receives salary and benefits. All other authors report there are no competing interests.

